# Reovirus infection induces transcriptome-wide unique A-to-I editing changes in the murine fibroblasts

**DOI:** 10.1101/2024.02.10.579760

**Authors:** Ayesha Tariq, Helen Piontkivska

## Abstract

The conversion of Adenosine (A) to Inosine (I), by Adenosine Deaminases Acting on RNA or ADARs, is an essential post-transcriptional modification that contributes to proteome diversity and regulation in metazoans including humans. In addition to its transcriptome-regulating role, ADARs also play a major part in immune response to viral infection, where an interferon response activates interferon-stimulated genes, such as ADARp150, in turn dynamically regulating host-virus interactions. A previous report has shown that infection from reoviruses, despite strong activation of ADARp150, does not influence the editing of some of the major known editing targets, while likely editing others, suggesting a potentially nuanced editing pattern that may depend on different factors. However, the results were based on a handful of selected editing sites and did not cover the entire transcriptome. Thus, to determine whether and how reovirus infection specifically affects host ADAR editing patterns, we analyzed a publicly available deep-sequenced RNA-seq dataset, from murine fibroblasts infected with wild-type and mutant reovirus strains that allowed us to examine changes in editing patterns on a transcriptome-wide scale. To the best of our knowledge, this is the first transcriptome-wide report on host editing changes after reovirus infection. Our results demonstrate that reovirus infection induces unique nuanced editing changes in the host, including introducing sites uniquely edited in infected samples. Genes with edited sites are overrepresented in pathways related to immune regulation, cellular signaling, metabolism, and growth. Moreover, a shift in editing targets has also been observed, where the same genes are edited in infection and control conditions but at different sites, or where the editing rate is increased for some and decreased for other differential targets, supporting the hypothesis of dynamic and condition-specific editing by ADARs.

---

Reoviruses, first isolated from human stool (Sabin, 1959) are non-enveloped dsRNA viruses (Forrest and Dermody, 2003). The viral genome is housed inside an icosahedral core, consisting of 10 segments classified as sigma (σ), mu (μ), and lambda (λ) based on their molecular weight (Danthi et al., 2013). Each RNA segment except for the one of four small segments encodes for a single protein (Eledge et al., 2019). Three genetically distinct but morphologically similar groups of reoviruses, type 1 Lang (T1L), type 2 Jones (T2J), and type 3 Dearing (T3D), are described based on nucleotide differences in their σ1 attachment protein (Douville et al., 2008), of which T3D has been described to be oncolytic (Gauvin et al., 2013). They are frequently found in untreated sewage and can infect different mammalian species, including humans and mice (Douville et al., 2008), through fecal, oral, and airborne routes (DeBiasi and Tyler, 2015). Reovirus infection most commonly occurs in infants and young children (Pruijssers et al., 2013) and shows respiratory, gastrointestinal, and central nervous system (CNS) manifestations (DeBiasi and Tyler, 2015). Although reoviruses are known to cause mostly mild disease in humans, viral strains have been isolated from children with severe meningitis and encephalitis, indicating neurovirulence (Ouattara et al., 2011). A novel strain isolated in Japan from children with meningitis and gastroenteritis, was found to be genetically similar to a strain identified from bats in China, suggesting potential interspecies transmission (Yamamoto et al., 2020).

The effectiveness of infection differs among 3 strains depending upon the route of infection (Gauvin et al., 2013) and the age of the host (Wu et al., 2018). For instance, the T3D strain causes lethal encephalitis in young mice with no distinct signs of infection in older ones (Wu et al., 2018).

Both T1 and T3D types can cause infection of CNS in mice. Differences in viral tropism arise due to the different sites of attachment for viral entry (Danthi et al., 2013). Although the site of attachment is strain-specific, reoviruses generally gain entry into the cell through endocytosis. Once inside the cell, viral RNA is sensed by the host’s cytosolic receptors (Pruijssers et al., 2013) leading to the induction of type I interferon response and ultimately expression of interferon-stimulated genes (ISGs) to stop viral replication (Stanifer et al., 2017). Reovirus infection also elicits type III interferon response, although the mechanism remains poorly understood (Abad and Danthi, 2020).

Replication of reovirus inside the host cell is inhibited by interferon-inducible protein-dependent kinase or PKR, and reoviral protein σ3 competes with PKR for binding to dsRNA and confers a partial resistance to PKR induced by type 1 IFN response (Rudd and Lemay, 2005). Cancerous cells have altered IFN response and decreased PKR activity, hence, reoviruses can preferentially replicate and survive (Phillips et al., 2018). A reovirus strain P4L-12 generated through chemical mutagenesis of wild-type T3D shows hypersensitivity to interferons with a 1000-fold decrease in viral titer inside interferon-treated cells compared to wild-type (Rudd and Lemay, 2005). RNA capping enzyme λ2 of reoviruses adds a 5’cap structure on viral mRNA to mimic host mRNA and escape the innate immune response. Viral strains with mutant λ2 are more sensitive to interferon-induced restriction by innate immunity (Hyde and Diamond, 2015). P4L-12 is a preferred oncolytic agent and has one amino acid substitution in RNA capping enzyme λ2 at position T636M, which is shown to be responsible for the difference in interferon sensitivity between T3D and P4L-12 (Sandekian and Lemay, 2015).

Infection of wild-type reovirus (ReoV) induces a strong IFN response (Hood et al. 2014; Pfaller et al., 2021) activating Adenosine Deaminases Acting on RAN (ADARs) as a downstream consequence of the IFN activation pathways (Pfaller et al. 2021). ADARp150, a long isoform of ADAR1, is an ISG and is present in both the cytoplasm and nucleus while the short isoform p110 is predominantly localized in the nucleus (Pfaller et al., 2021). ADAR1 is known to edit viral RNA, rendering it destabilized for replication. However, the editing is dynamic and can be pro- or antiviral depending on the type of virus (Piontkivska et al., 2021, Tang et al., 2015). Studies from ADAR1/2 knockouts found no impact on ReoV replication (Ward et al., 2011), potentially because of the protective core-like sub-virion particle that shields viral genomes during replication (Pfaller et al. 2021). Studies of animal reoviruses, such as that of grass carp reovirus (GCRV), showed activation of an ADAR1 homolog in response to viral infection (Yang et al., 2012; Rao and Su 2015), implying a role of the enzyme in the antiviral response, although its impact on the viral editing remains to be elucidated.

It has been established that RNA viruses, including reoviruses, experience ADAR editing during infection (eg, Tang et al., 2015, Pfaller et al., 2021, Piontkivska et al., 2021, Ringlander et al., 2022, Zhu et al., 2023). However, the impact of viral infection on the editing of the host transcriptome is not well understood (Piontkivska et al., 2021). A study from neonatal mice injected with reovirus T3D into the brain found that despite strong induction of ADARp150, the majority of surveyed editing substrates exhibited little to no editing changes, including no detectable changes within *5HT2C* (serotonin 2C receptor) transcripts (Hood et al., 2014). Nonetheless, some genes did experience changes in editing, such as cytoplasmic FMR1 interacting protein 2 (*Cyfip2*), filamin A (*Flna*), and bladder cancer-associated protein (*Blcap*) (Hood et al. 2014), indicating that infection-induced changes in editing patterns may be nuanced and possibly influenced by the stage of development. To study the effect of reovirus infection on ADAR editing of the host at a whole transcriptome scale, we took advantage of a publicly available deep-sequenced BioProject PRJNA320288 dataset, from Boudreault et al., 2016 (GEO accession number GSE81017) where the L929 mice cell lines were infected with wild-type and mutant ReoV strains, and compared those to the uninfected control cells (Table 1).

**Table 1:**
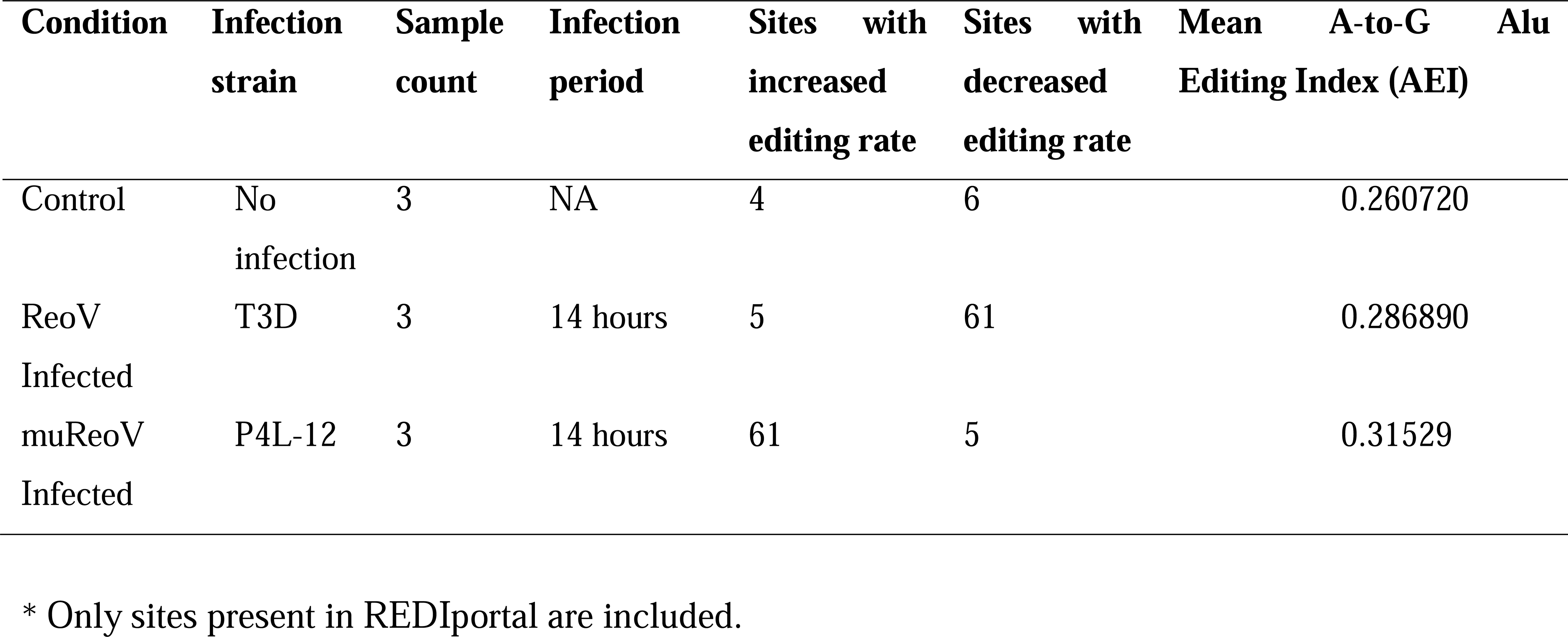
Characteristics of the BioProject (PRJNA320288) dataset used in the study. ReoV infected represents fibroblasts (L929) cells infected with wild-type reovirus (ReoV) T3D, while muReoV denotes cells infected with mutant reovirus (muReoV) P4L-12. Total RNA was extracted 14 hours after infection (Boudreault et al., 2016). Edited sites* with increased or decreased editing were calculated as control vs infection, and ReoV infection vs muReoV infection, respectively.

The Automated Isoform Diversity Detector (AIDD) pipeline (Plonski et al., 2020) was used to map the reads, infer ADAR expression, and predict ADAR editing sites (Supplemental_Table_1). DESeq2 R package (Love et al., 2014) was used for differential expression analysis of ADAR genes and transcripts. The Alu Editing Index (AEI) as a global measure of hyperediting was calculated per sample using the RNAEditingIndexer tool (Roth et al., 2019). Variant frequency counts estimated using bam-readcount (Khanna et al., 2022) was used to calculate the average editing rate per site. The editing rate was defined as the number of G reads for an A reference site or C reads for a U(T) reference site on an opposite strand, divided by the total number of reads mapped to that site. Average editing was calculated by taking the mean of editing rate for a single site across all samples for a specific condition. Control samples were compared with infected samples for differential editing changes. A similar comparison was made for samples infected with wild-type vs mutant strains. To ensure that only confirmed editing sites are considered, only those editing sites with reference in the REDIportal, a comprehensive nonredundant database of A-to-I editing events (Picardi et al., 2017), were included. Other filtering options required that the edited site had greater than or equal to 20 total reads aligned, and that editing rates were greater than 0.01, or less than 0.99, and not between 0.49 and 0.51, to remove potential noise, homozygous genomic variants, and heterozygous genomic variants, respectively. Pathway enrichment analysis was done using The Database for Annotation, Visualization, and Integrated Discovery (DAVID) functional annotation tool (Huang et al., 2009, Sherman et al., 2022). R package biomaRt (Durinck et al., 2009) was used for annotation of differentially expressed genes. We used the Ensembl Variant Effect Predictor (VEP) to analyze the potential impact of novel A-to-I edits on the transcript (McLaren et al., 2016).

The expression of ADAR1, in transcript per million (TPM), was higher in infected samples than in control (ANOVA, p = 2.23E-05). While the TPM counts for ADAR2 and ADAR3 were relatively low, nonetheless, they also differed significantly (ANOVA, p=8.34E-05, and p=0.01633633, respectively) among three conditions (Figure 1A, Supplemental_Table_2). Differential expression analysis based on transcript counts using DESeq2 confirmed these findings, where the expression of ADARp110 and ADARp150 transcripts was significantly higher, more than two-folds, in both ReoV- and muReoV-infected samples compared to controls, with padj values < 0.05 (Supplemental_Table_3 & Supplemental_Table_4). However, at the gene level only ADAR1, but not ADAR2 or ADAR3, was among differentially expressed genes (DEGs) (Supplemental_Table_4A) for the ReoV-vs-control comparison, with log2FoldChange at ∼2.48 (padj values < 0.05), while ADAR2 was among DEGs for the muReoV-vs-control comparisons, respectively, although the fold change was much smaller, at ∼ 1.12 (padj values < 0.05) (Supplemental_Table_4C).

**Figure 1:**
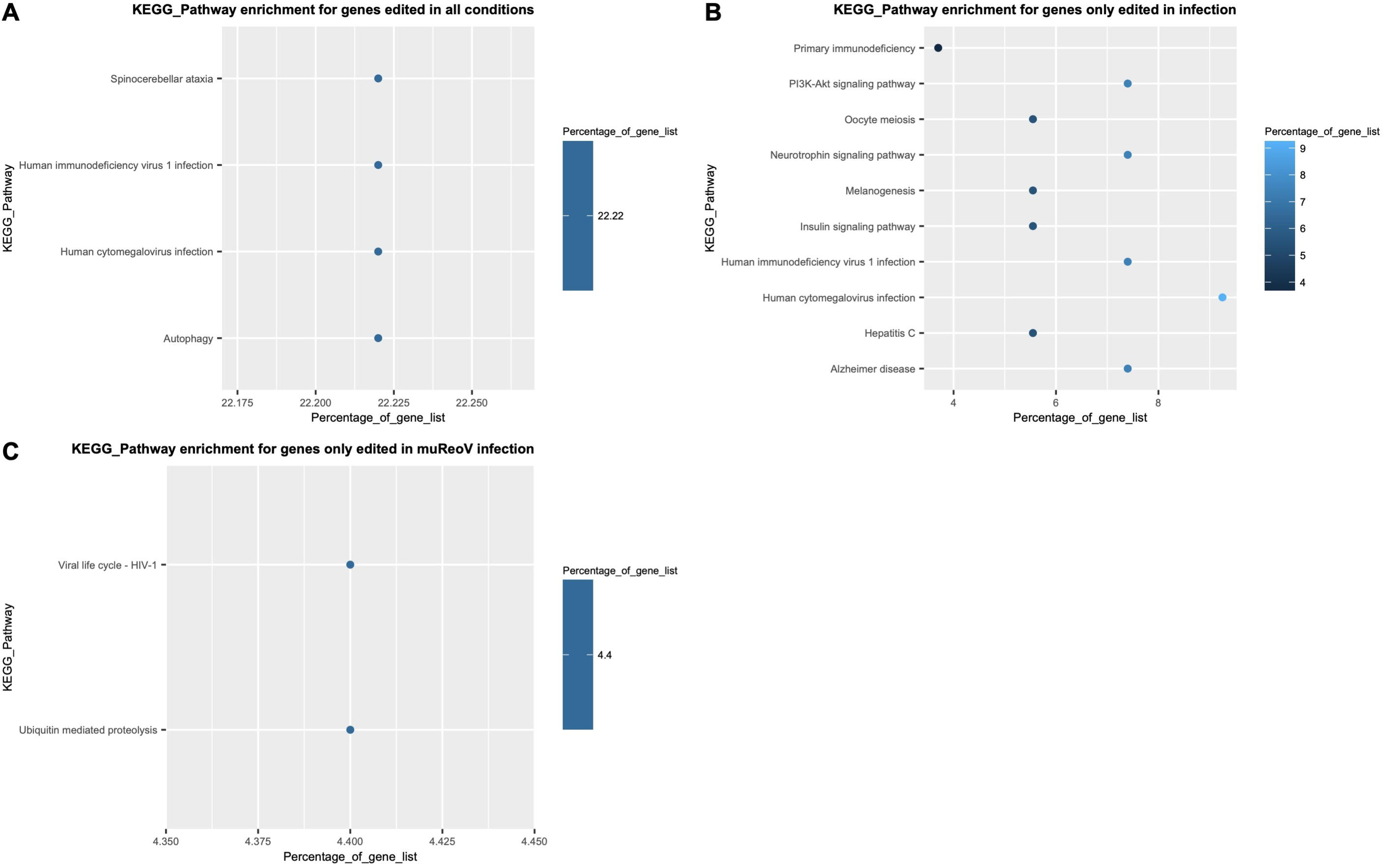
Differences in expression of ADAR genes, and distribution of ADAR editing between ReoV-infected and uninfected control samples. Panel A shows the average TPMs (transcripts per million) for ADAR1, ADAR2 and ADAR3 across all 9 samples. Detailed DESeq2 results based on gene and transcript counts support statistically significant differences in ADAR expression, with padj < 0.05 (Supplemental_Table_4). Panel B shows the number of region-specific editing changes with an editing rate > 0.01 in all samples. Panel C shows a comparison of the Alu Editing Index (AEI) for all three conditions. The A-to-G editing index is higher in infected samples. Panel D shows mean editing rate per site per sample for all three conditions. More sites are edited in infected samples, with highest editing rate in muReoV infection.

Next, we examined region-specific editing and differential editing per site. The highest number of edited sites falls in 3’UTRs, while 5’UTRs are the least edited, among all edited genomic regions (Figure 1B). The Alu Editing Index (AEI), for A-to-G changes, is also high for infected samples with C-to-U(T) being the next most common transition (Figure 1C, Supplemental_Table_5). The initial comparison, for all the sites including those that did not map to REDIportal genes, showed that there is a shift in editing between control, ReoV infected, and muReoV infected samples, where infected and mutant infected samples show an increase in the number of editing sites (Figure 1D). Overall, control, reovirus T3D or ReoV infected and mutant P4L-12 or muReoV infected samples had 487, 784, and 1284 A-to-G or U(T)-to-C edited positions respectively (Supplemental_Figure_1, Supplemental_Table_6).

Next, all the A-to-G or U(T)-to-C substituted sites were filtered for those mapped to REDIportal genes. Only 17 sites were mapped to genes from control samples while in wild-type (ReoV) and mutant reovirus-infected samples (muReoV), 71 and 153 sites were mapped to REDIportal genes respectively (Supplemental_Figure_2, Supplemental_Table_7). Overall, only 9 positions were in common between infection and control, with 6 positions shared across all three conditions, with significant differential editing for a few positions. Mean editing of common sites was higher in infection than in control, with the *PUM2* gene experiencing the highest amount of change (Figure 2, Supplemental_Table_8). Similarly, 66 sites among sites unique to infection samples were shared between ReoV and muReoV infection, and the overall editing rate was higher in muReoV samples, with the highest mean editing in the *Tapbp* gene (Table 2, Supplemental_Table_9). Moreover, 87 positions were only edited in mutant-infected samples (Supplemental_Table_10), while only 5 positions were unique to ReoV-infected samples (Supplemental_Table_11).

**Figure 2:**
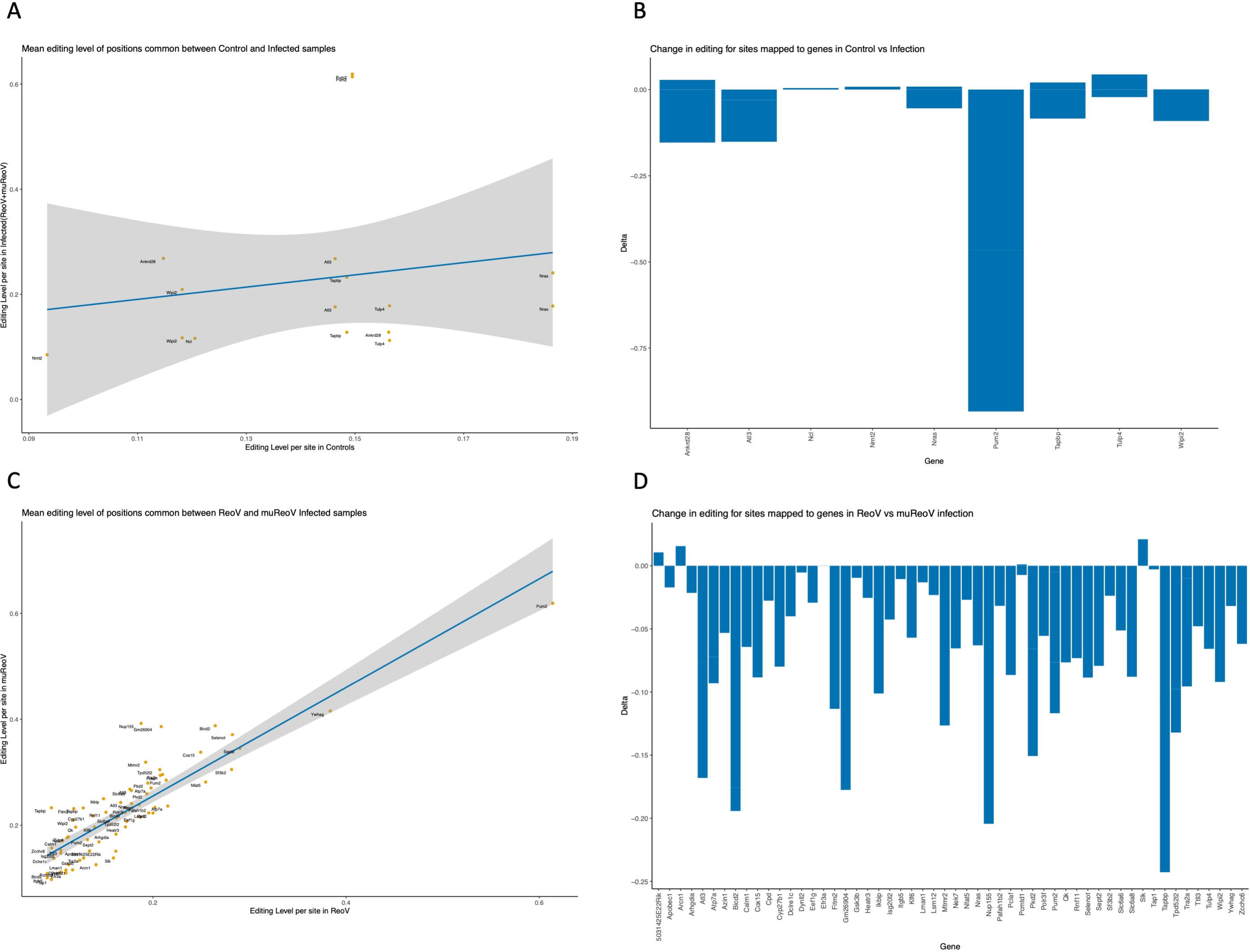
Difference in mean editing rate, per site, between control and infected samples. Only sites present in REDIportal are shown. Panel A represents mean editing for sites common across all control and infected samples. The grey area represents 95% CI for the difference in editing. Panel C shows mean editing for sites common in muReoV and ReoV only. Panels B and D show the difference in mean editing per site for control vs infection (ReoV + muReoV) and ReoV vs muReoV only respectively. The full lists of sites available in Supplemental_Table_8 and Supplemental_Table_9.

**Table 2:**
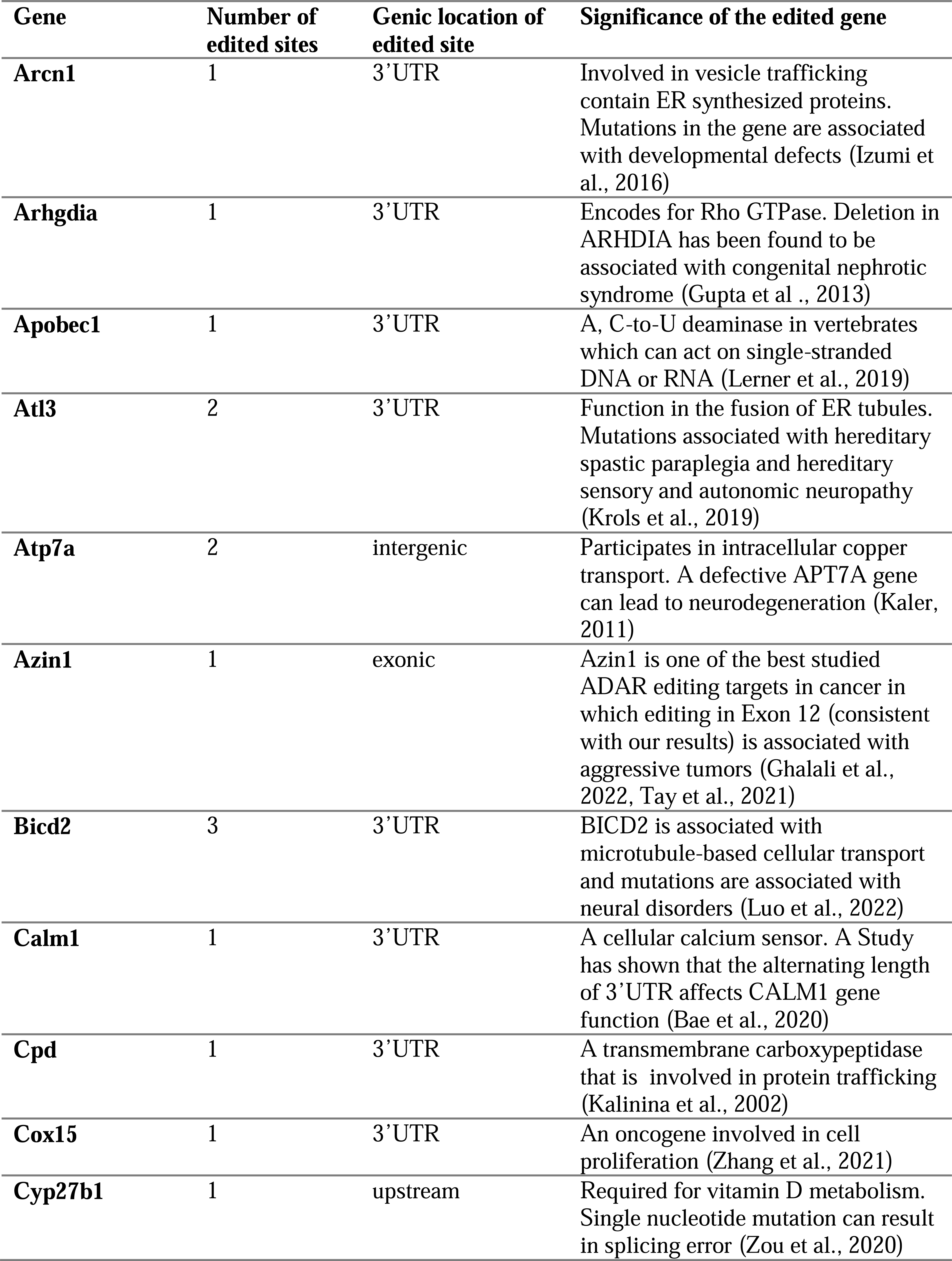
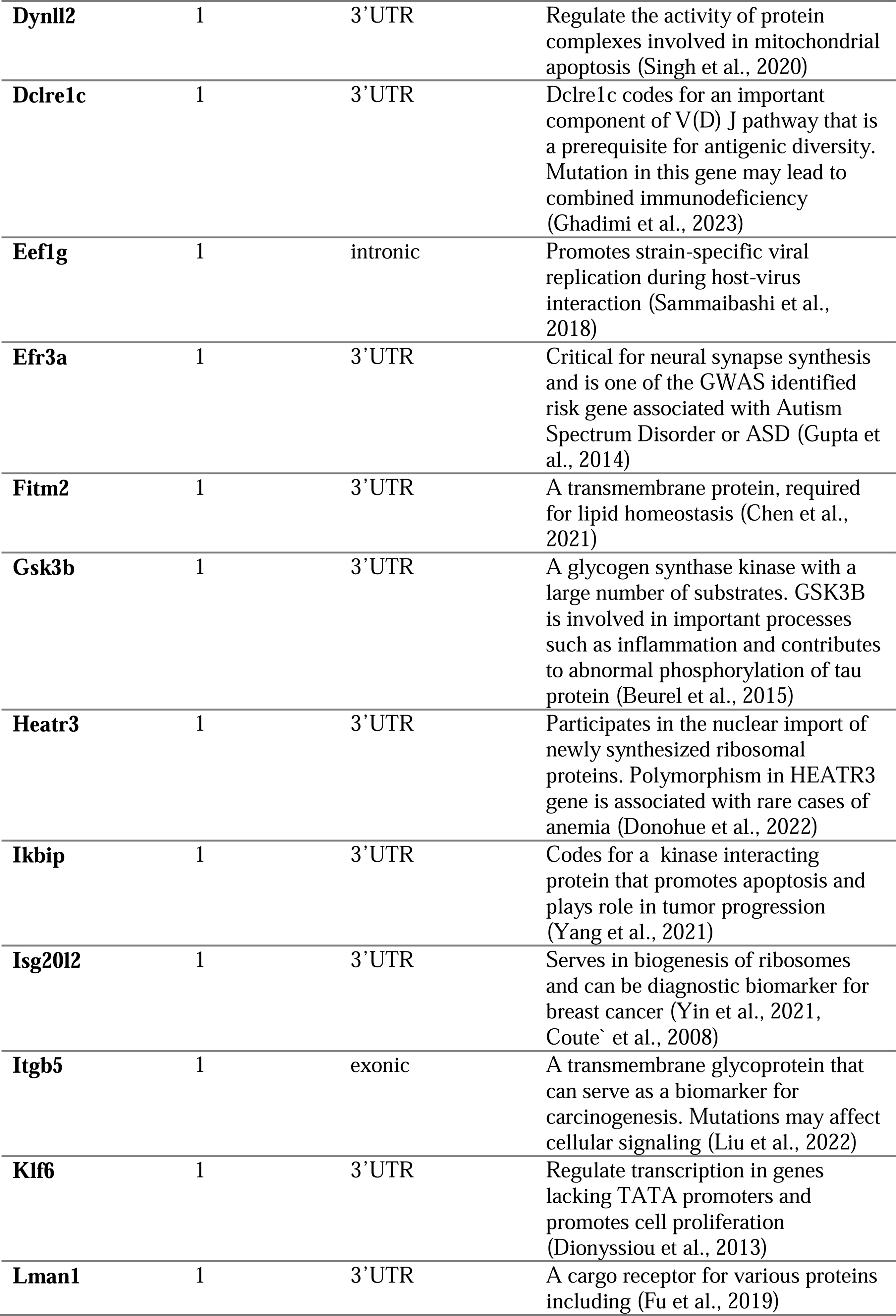
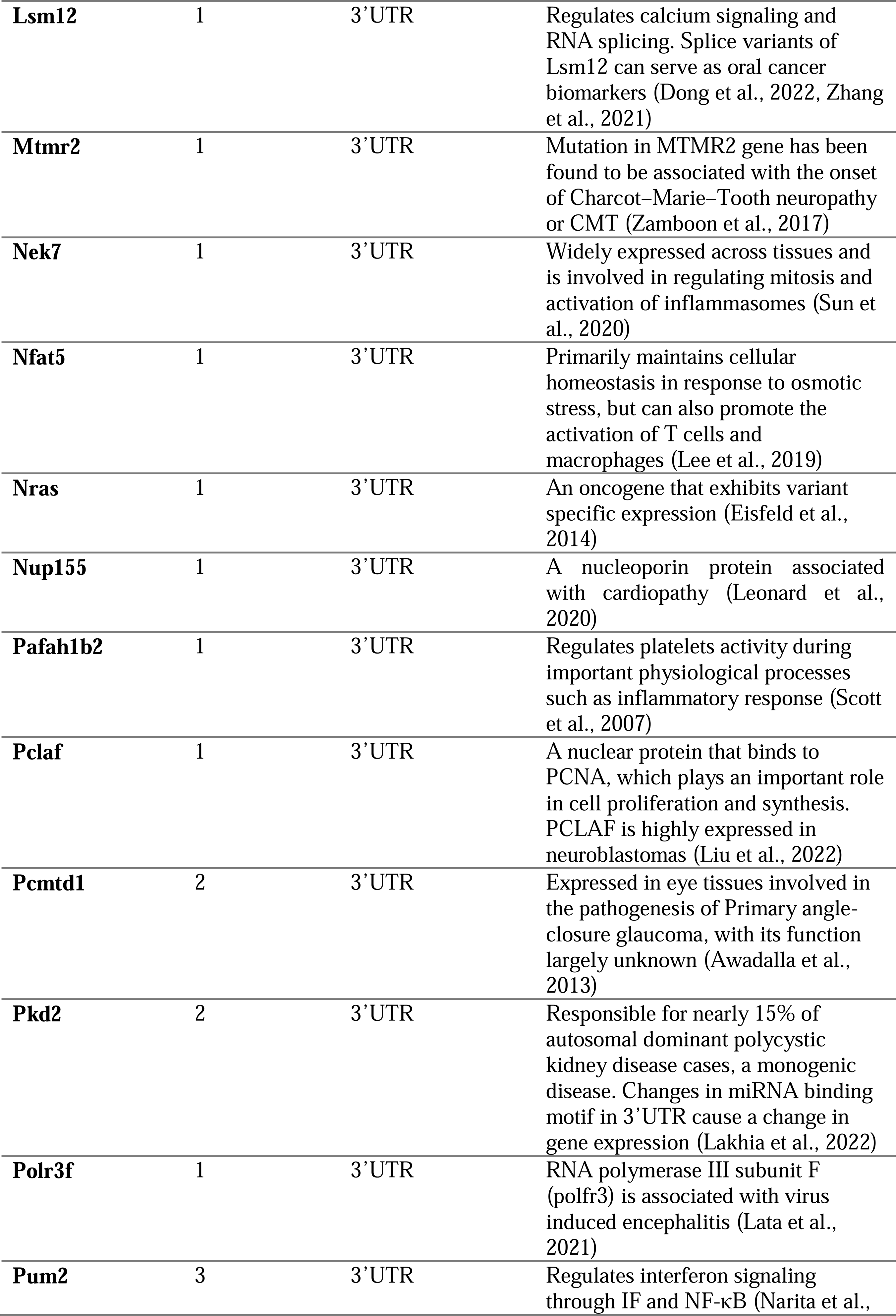
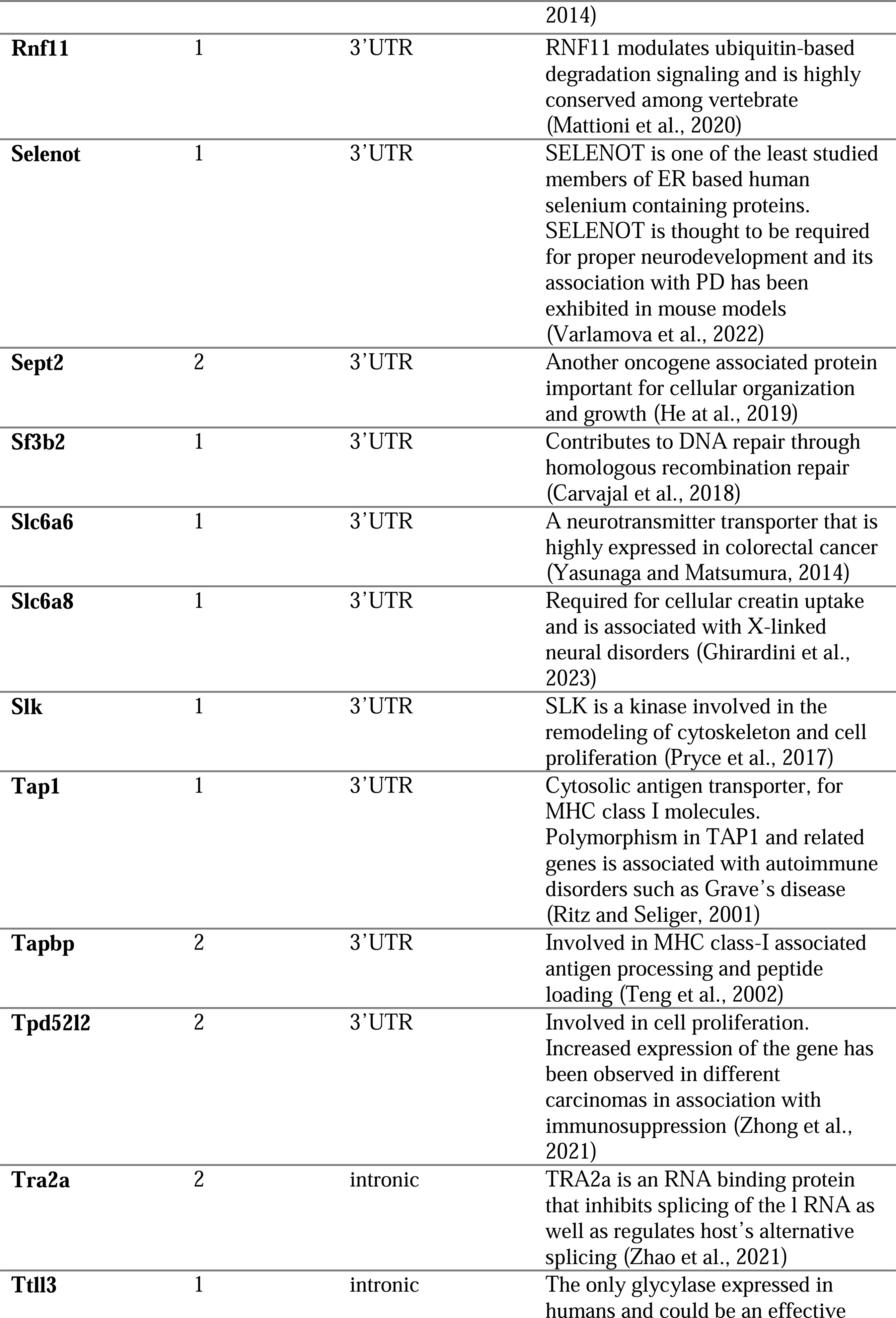
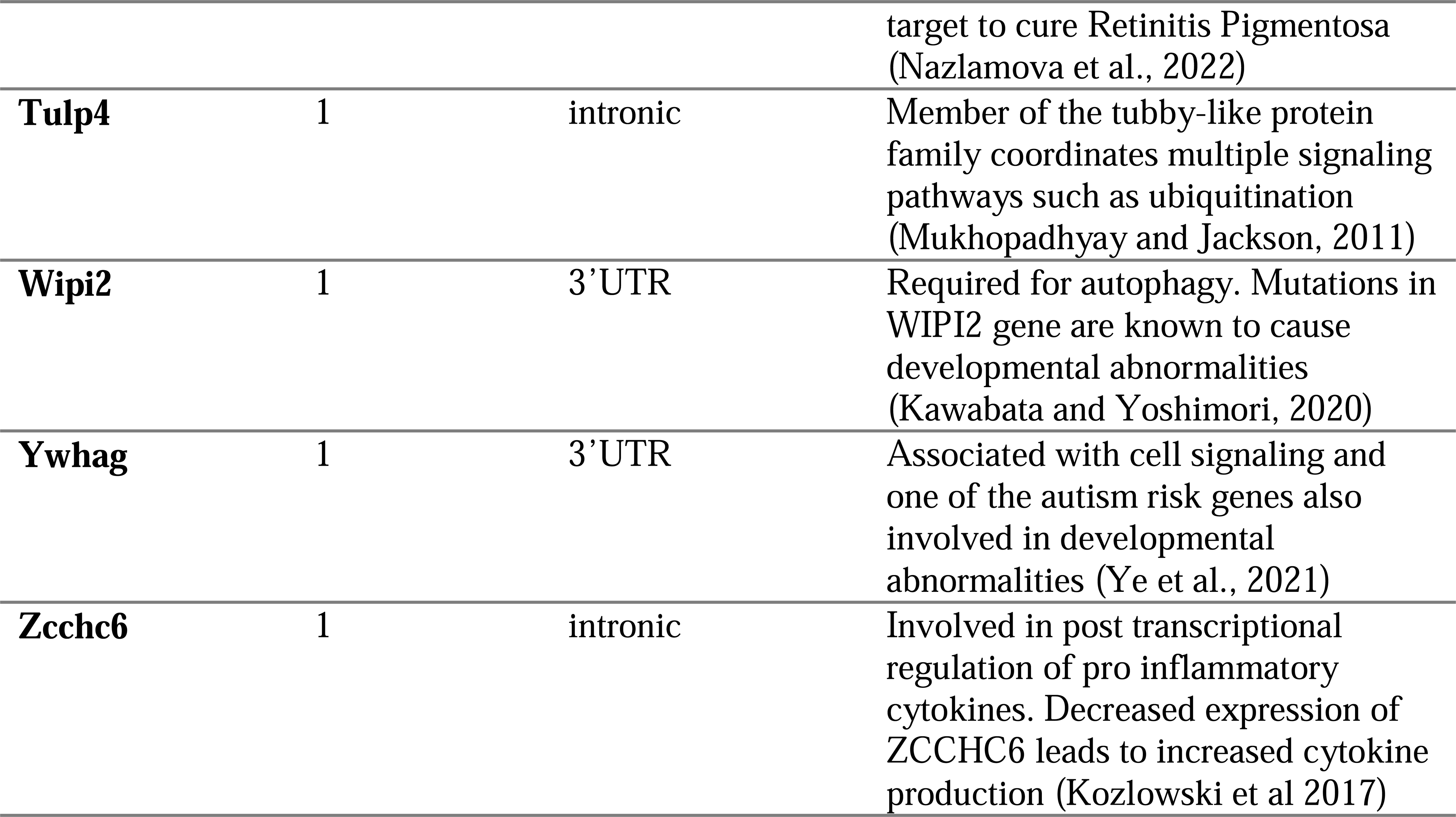
List of genes with significant ADAR editing sites that are unique to infection samples, along with the region of change, number of edited sites in each gene, and functional relevance of the gene.

Our results from pathway enrichment analysis show a shift in affected pathways from control to infection (Figure 3). Not only different pathways are affected in infected samples compared to control, but editing in infected samples also appears to be spread across a larger number of pathways than that of control samples. Gene-specific descriptions of expanded biological categories from InnateDB (Lynn et al., 2008, Lyn et al., 2010, Breuer et al., 2013) can be found in Supplemental_Table_12.

**Figure 3:**
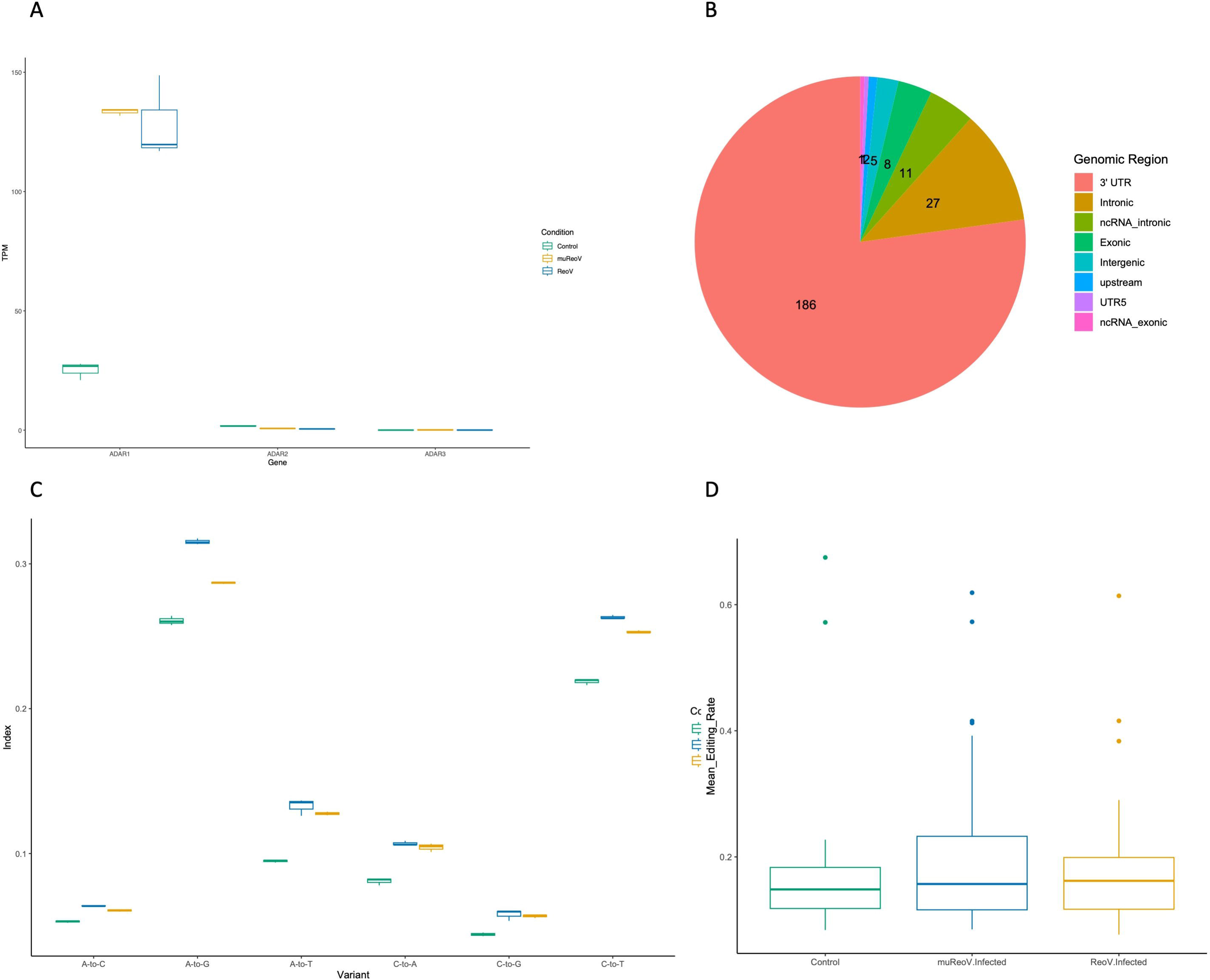
KEGG pathway enrichment for sites in all treatment conditions and unique to infection samples. Panels A, B and C show pathway enrichment for genes edited in all conditions (A), common in muReoV and ReoV (B), and unique to muReoV (C), respectively.

One of the many important components of the host-virus interaction dynamics is the editing of viral as well as host RNA by host ADARs (Piontkivska et al., 2021). Our results demonstrate that the reovirus infection induces nuanced changes in the ADAR editing pattern of the host transcriptome. Reovirus infection causes an increase in the ADAR1 expression that translates to editing changes that are not unidirectional but dynamic, where some sites may experience consistent increase in editing in infection, while other sites experience consistent decrease. Genes from our analysis that experienced ADAR editing participate in cellular metabolism, growth, and immune function pathways. Interestingly, editing mediated by infection with muReoV strain is shown to affect ubiquitin-mediated proteolysis (Figure 3C), which could be an indication of misregulation of translation, potentially benefiting the survival of the virus (Lou, 2016). We also have observed a shift in editing targets, where the same gene is being edited in control and infection, but at a different position. For instance, the Atl3 gene associated with cellular integrity and cytokinesis (Mohammadi et al., 2023) has only one edited site in 3’UTR at positions 7538115 and 7535531 in control and muReoV respectively. One explanation for that phenomenon could be that RNA editing is condition-specific (Liew et al., 2017), and/or reflect the effect of other unknown factors that regulate host-virus interactions (Piontkivska et al., 2021).

It is known that reoviruses can preferentially infect and replicate inside Ras-transformed cells, leading to oncolysis (Phillips et al., 2018). Reovirus infection also induces an anti-tumor immune response in cancerous cells, independent of activated Ras signaling, but the exact mechanism is not fully elucidated (Gong et al., 2016, Prestwich et al., 2008, Philips et al., 2018). It can be speculated that differential ADAR editing is one of the host factors contributing to reovirus-induced anti-tumor immunity. For instance, Pumilio proteins, *PUM1* and *PUM2*, are evolutionarily conserved proteins and have been identified as positive regulators of interferon signaling upon viral infection, by meditating the activation of interferon response factor (IF) and NF-κB (Narita et al., 2014). In our analysis of edited sites unique to infection samples, the *PUM2* gene was found to be one of the most edited genes (Table 2), with 3 edited positions in 3’UTR. Moreover, *Tapbp* gene (Table 2) was only edited in infection samples with increased editing in muReoV infection. *Tapbp* (TAP binding protein) mediates the interaction between MHC and TAP molecules to effectively load and transport antigenic peptides (Boyle et al., 2012). These results suggest that the editing of host transcriptome by ADARs can potentially affect cellular signaling and immune processes. The higher average expression of ADAR1 in the muReoV infection compared to that of wild-type infected samples indicates that even low viral load, as expected in interferon-sensitive strain, can induce high ADAR1 expression that translates to differential editing of host transcriptome. Editing patterns may vary depending upon not only viral but also host genetics (Gelbart et al., 2020). Studies have also pointed out that the oncolytic action of reovirus can differ based on cell type and tumor microenvironment (Philips et al., 2018, Alain et al., 2007).

Our analysis of transcriptome-wide RNA editing changes in murine fibroblasts conclusively shows that infection with reoviruses indeed affects the global editing landscape, leading to site-specific unique editing changes. Similar to prior studies, majority of editing changes observed here are located within 3’UTRs (e.g., Wales-McGrath et al., 2023), underscoring the need to understand the consequences of such seemingly minor and likely dynamic changes on regulation of gene expression. Variants in 3’UTRs can influence gene expression through interruption of micro-RNA binding, polyadenylation, and posttranscriptional processing (Skeeles et al., 2013); changes in UTRs can also contribute to disease progression by affecting mRNA expression (Hong and Jeong, 2023).

The effect of viral infections on the RNA editing patterns of the host remains largely underexplored, with very little if any data available for many viral families (Piontkivska et al., 2021). Future ReoV studies should focus on tissue and tumor-specific patterns of ADAR editing in humans, for a safer reovirus-based oncolytic therapy. Comparison of the host’s editing landscape after reovirus infection with related pathogens, such as rotavirus (Crawford et al., 2017), could clarify whether or how ADAR editing is affected by the severity of infection. In our analysis, we also identified 2314 novel A-to-G or U(T)-to-C changes (Supplemental_Table_13) that were not reported in REDIportal. Many of these sites fall in intergenic regions and may have modifier effects on the transcript (Supplemental_File_14). Experimental validation of these sites can offer further insights into the effect of reovirus infection on host transcriptome. While this study is limited by a small sample size, future studies should offer larger samples and include diverse tissue/cell types, to expand our understanding of the interaction of reoviruses with host cells.

## Supplemental files and legends

Supplemental_Table_1: Read depth for all samples.

Supplemental_Table_2: Average TPM values for ADAR1,2 & 3 in all samples.

Supplemental_Table_3: DESeq2 differential ADAR transcript expression results for ADARp150 and ADARp110.

Supplemental_Table_4: DESeq2 gene and transcript expression analysis results for all edited genes.

Supplemental_Table_5: Alu/SINE Editing Index (AEI).

Supplemental_Table_6: All edited sites in Control and Infected samples with >20 reads aligned and editing rate > 0.01.

Supplemental_Table_7: All edited sites in Control and Infected samples with >20 reads aligned and editing rate > 0.01 and annotated from REDIportal.

Supplemental_Table_8: Edited sites common between control, ReoV and muReoV.

Supplemental_Table_9: Edited sites common between ReoV and muReoV only.

Supplemental_Table_10: Edited sites unique to muReoV-infected samples.

Supplemental_Table_11: Edited sites unique to ReoV-infected samples.

Supplemental_Table_12: Pathway enrichment results from InnateDB for genes uniquely edited in infection.

Supplemental_Table_13: Novel edited sites in all conditions that were not documented in REDIportal.

Supplemental_File_14: Impact of Novel edited sites on transcriptome. These results are compiled from Ensembl Variant Effect Predictor (VEP) tool, for each treatment condition.

Supplemental_Figure_1: Distribution of ADAR edited sites among control and reovirus-infected, with reovirus T3D, or ReoV, and mutant P4L-12, or muReoV, infected samples. There was a total of 487, 784, and 1284 A-to-G or U(T)-to-C edited positions across three treatment categories. The full list is available in Supplemental_Table_6.

Supplemental_Figure_2: Distribution of a subset of ADAR edited sites that are also present in REDIportal genes among control and reovirus-infected, with ReoV or muReoV, samples. There was a total of 17, 71 and 153 A-to-G or U(T)-to-C edited positions across three treatment categories. The full list is available in Supplemental_Table_7.

## Supporting information

Supplemental_Tables_1-through-13

## List of abbreviations

ADAR: Adenosine Deaminases Acting on RNA
ReoV: Reovirus
muReoV: Mutant Reovirus
UTR: Untranslated Region
ISG: Interferon Stimulated Gene
PKR: Protein Kinase R
CI: Confidence Interval

