## Supplementary figures and images for "Reovirus infection induces transcriptome-wide unique A-to-I editing changes in the murine fibroblasts"

### Supplemental_Fig_1

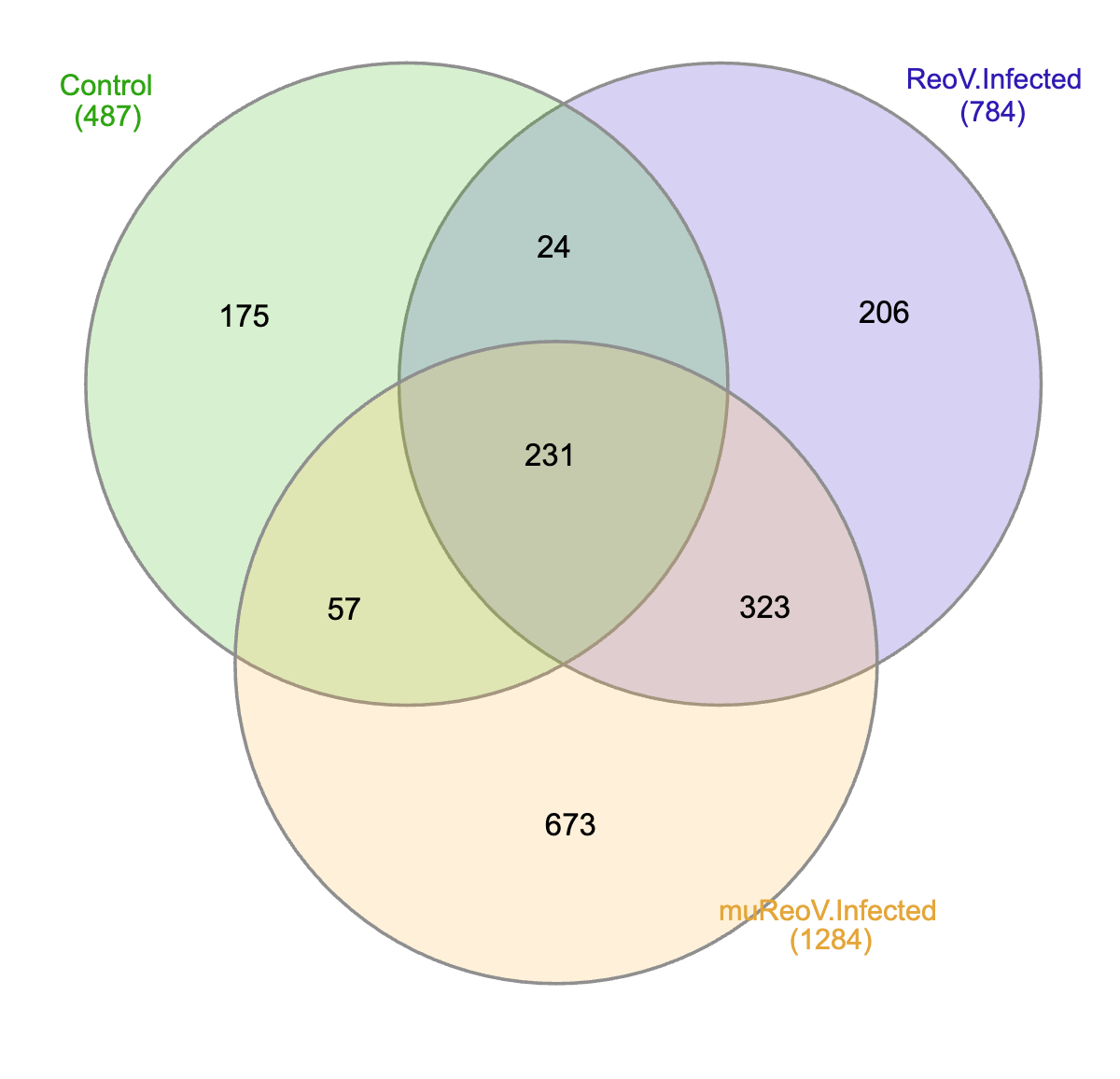

### Supplemental_Fig_2

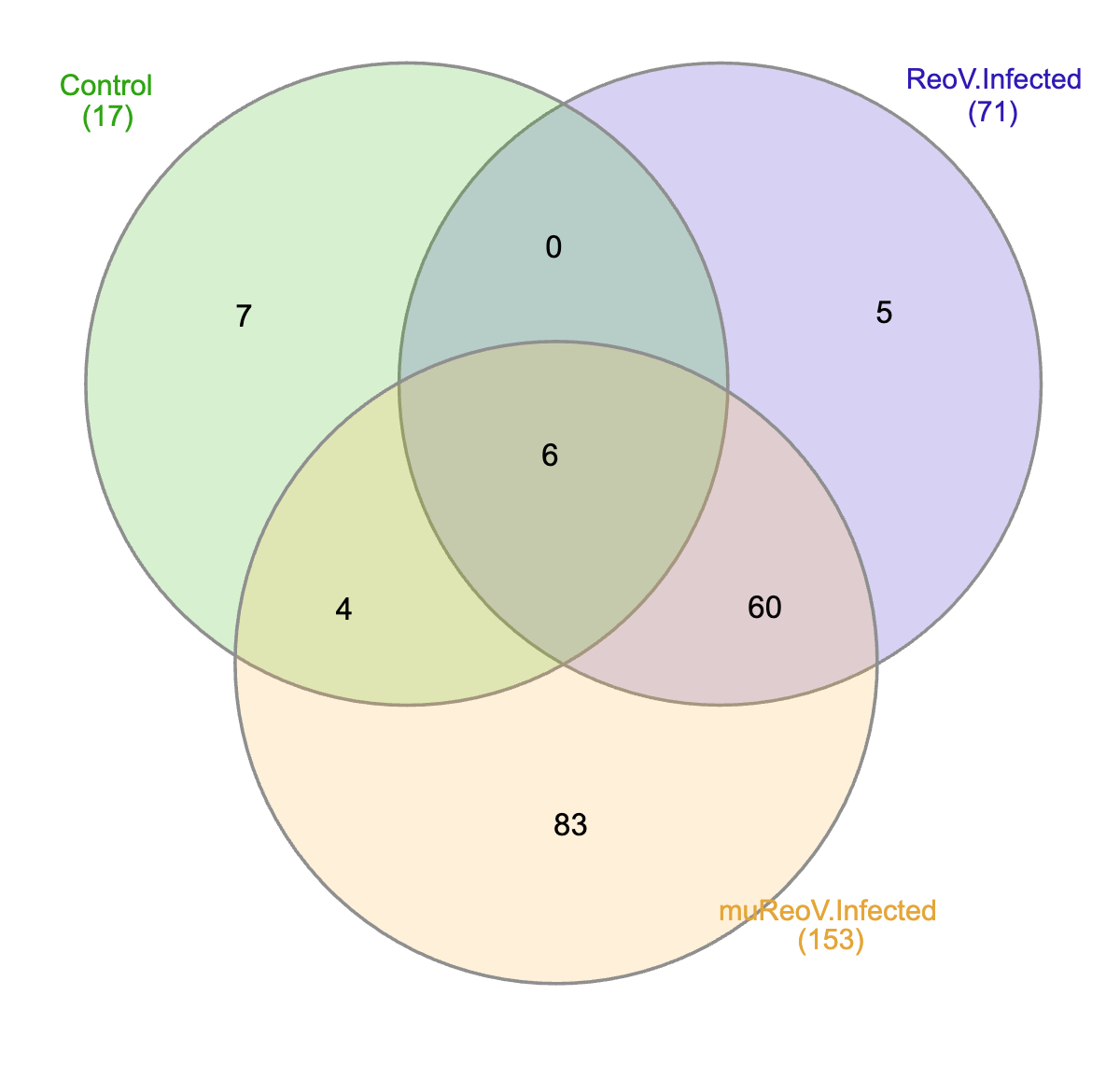
